# No behavioral evidence for rhythmic facilitation of perceptual discrimination

**DOI:** 10.1101/2020.12.10.418947

**Authors:** Wy Ming Lin, Djamari A. Oetringer, Iske Bakker-Marshall, Jill Emmerzaal, Anna Wilsch, Hesham A. ElShafei, Elie Rassi, Saskia Haegens

## Abstract

It has been hypothesized that internal oscillations can synchronize (i.e., entrain) to external environmental rhythms, thereby facilitating perception and behavior. To date, evidence for the link between the phase of neural oscillations and behavior has been scarce and contradictory; moreover, it remains an open question whether the brain can use this tentative mechanism for active temporal prediction. In our present study, we conducted a series of auditory pitch discrimination tasks with 181 healthy participants in an effort to shed light on the proposed behavioral benefits of rhythmic cueing and entrainment. In the three versions of our task, we observed no perceptual benefit of purported entrainment: targets occurring in-phase with a rhythmic cue provided no perceptual benefits in terms of discrimination accuracy or reaction time when compared with targets occurring out-of-phase or targets occurring randomly, nor did we find performance differences for targets preceded by rhythmic vs. random cues. However, we found a surprising effect of cueing frequency on reaction time, in which participants showed faster responses to cue rhythms presented at higher frequencies. We therefore provide no evidence of entrainment, but instead a tentative effect of covert active sensing in which a faster external rhythm leads to a faster communication rate between motor and sensory cortices, allowing for sensory inputs to be sampled earlier in time.

## Introduction

When presented with rhythmic input we tend to produce rhythmic behavior. Think of clapping to a drumbeat and being able to continue clapping to the beat after the drum stops. Such rhythmic behavior, driven by temporal expectations, could be subserved by rhythmic brain activity (i.e., neural oscillations), a prominent feature of brain dynamics. In this view, internal neural oscillations that synchronize (entrain) to external environmental rhythms reflect temporal predictions, thereby facilitating perception and behavior (Lakatos et al., 2008).

This entrainment proposal rests on the key idea that neural oscillations reflect alternating excitability states (excitation/inhibition) of neuronal ensembles (Başar et al., 2013; Bishop, 1932). While there is some evidence that the phase of ongoing oscillations at the time of sensory stimulation impacts the neural response to that stimulus, as well as subsequent behavioral performance (e.g. Busch et al., 2009; Mathewson et al., 2009; Ten Oever & Sack, 2019; for a review see VanRullen, 2016), this evidence is far from conclusive as several studies have reported null results (e.g. Benwell et al., 2017; O’Hare, 1954; Ruzzoli et al., 2019; Vigué-Guix et al., 2020; Walsh, 1952).

Recently, the proposal of entrainment as a key mechanism for synchronizing with external input in order to optimize perceptual processing has gained traction, particularly in the fields of speech and language comprehension (for reviews see: Haegens & Golumbic, 2018; Meyer et al., 2019; Obleser & Kayser, 2019). However, there seems to be no consensus as to the definition of neural entrainment as a biophysical process (Haegens, 2020; Haegens & Golumbic, 2018; Lakatos et al., 2019; Obleser & Kayser, 2019). One such proposal (Haegens & Golumbic, 2018) — on which the current study is theoretically framed — argues for a strict definition of entrainment where: (1) an endogenous oscillator exists in the absence of rhythmic stimulation; that is, there is internally generated oscillatory brain activity at a certain frequency, (2) the endogenous oscillator adjusts its phase to align with external rhythmic stimulation, but only as long as the external rhythm falls within a range near that of the intrinsic frequency, and (3) the oscillatory activity continues for a number of cycles after the external rhythm stops.

Entrainment is often investigated with rhythmic cueing paradigms where participants are presented with a stimulus stream at a certain frequency. This rhythmic stream is then followed by a target stimulus that might occur in-or out-of-phase with the rhythmic cue, one or more cycles later (Jones et al., 2002). While several studies have shown that rhythmic cues indeed facilitate target processing, particularly for targets occurring in-phase (Jones et al., 2002, 2006; Rohenkohl & Nobre, 2011; Rohenkohl et al., 2011), others have reported opposite (Barnes & Johnston, 2010; Bauer et al., 2015; Hickok et al., 2015; Spaak et al., 2014, see Haegens & Golumbic for review) or null effects (Bosker & Kösem, 2017).

If entrainment indeed optimizes perception, we expect rhythmic cueing paradigms to produce certain behavioral benefits that follow from the criteria outlined above. Namely, we expect participants to perform better in conditions where temporal expectations are more readily built up: (1) when the cue is rhythmic (vs. random, arrhythmic, or continuous), i.e., providing explicit rhythmic temporal information, (2) when the target timing is rhythmically aligned with the cue (vs. occurring at a random time), i.e., providing implicit rhythmic structure, and (3) occurs in-phase (vs. out-of-phase) with respect to the cue. Further, we expect this behavioral benefit to wane over time as the entrained oscillation persists for a number of cycles after the external rhythm stops. Thus, we expect (4) participants to perform better for targets occurring shortly after the rhythm (vs. later). Finally, we expect (5) this behavioral benefit to be tightly linked to the frequency of the external rhythm, i.e., frequencies closest to endogenous oscillations are more behaviorally beneficial than others.

In a series of three behavioral experiments, we aimed to test these key predictions. A total of 181 healthy human participants performed an auditory pitch discrimination task where a target tone was presented after a rhythmic or random (i.e., continuous) auditory cue, with the timing of the target either rhythmically aligned to the cue or randomly timed. We manipulated the timings such as to be able to test each of our predictions listed above, and report no support for any of them.

## Methods

### Participants

Thirty-two healthy participants (21 female, 11 male; age range: 18–31 years, *median* = 23 years) took part in Experiment I. We excluded two participants from the analysis due to low performance levels and one participant due to low number of trials left after preprocessing. A total of 119 healthy participants (77 female, 42 male; age range: 18-35 years; *median* = 22 years) took part in Experiment II. Five participants were excluded due to low performance levels. Of the remaining participants, 30 performed the rhythmic cue-rhythmic target condition, 29 the rhythmic cue-random target condition, 29 the random cue-rhythmic target condition, and 26 the random cue-random target condition. Thirty healthy participants (22 female, 8 male; age range: 18–33 years, *median* = 22 years) took part in Experiment III. One participant was excluded due to low trial number. All participants provided written informed consent before testing and were fully debriefed about the goals of the study. The study was approved by the local ethics committee (CMO Arnhem-Nijmegen).

### Experimental task and stimuli

Participants performed an auditory target discrimination task in which they had to indicate whether a brief target tone either increased or decreased in pitch (Wilsch et al., 2020; Figure 1). The target tone consisted of 30 base frequencies that were randomly drawn from 500 to 1500 Hz. We modulated the pitch to either decrease or increase over time. This modulation was adjusted for each participant individually during a practice session, with a bigger slope (i.e., a larger difference between the pitch frequency at the start and end of the stimulus) being easier, such that participants performed at approx. 75% accuracy. The tone started and ended with a 10 ms cosine ramp fading in or out. The resulting target tone had a sample rate of 44100 Hz and a duration of 40 ms.

**Fig. 1:**
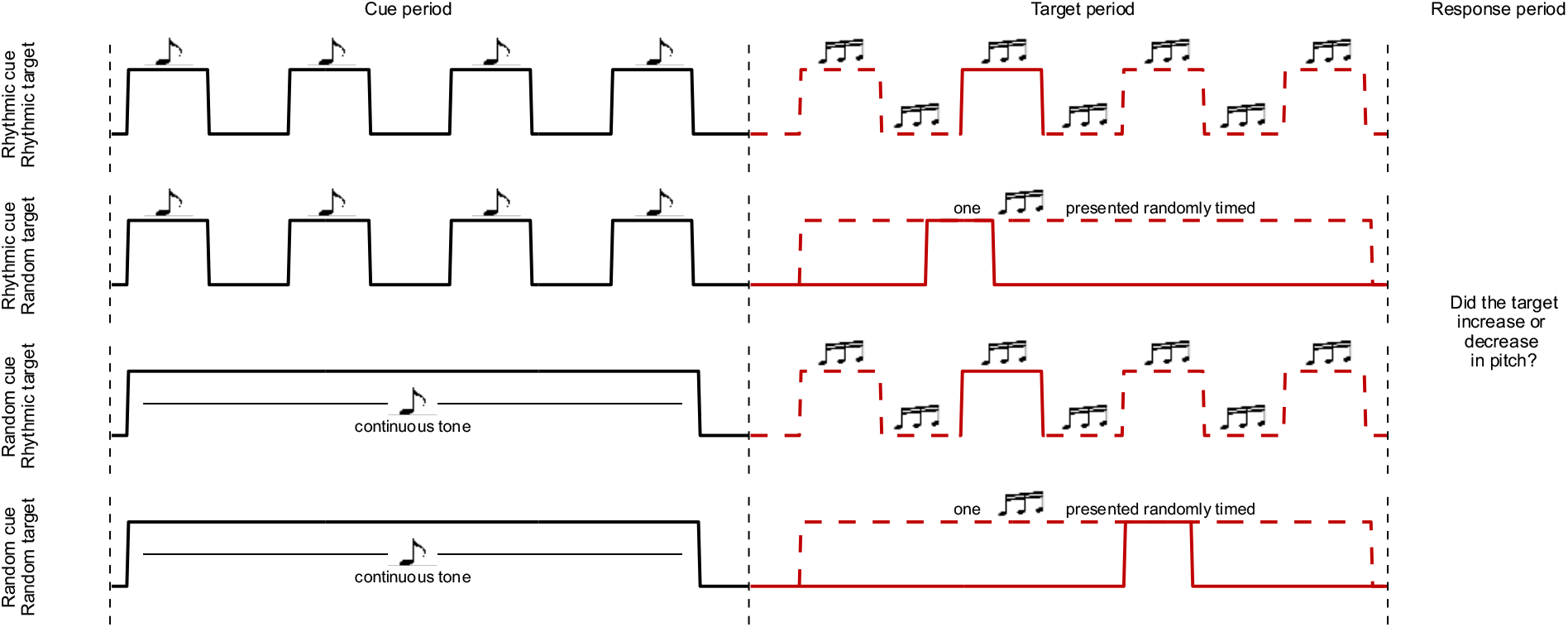
Experimental paradigm. All experiments used a variation of the auditory pitch discrimination task where a four-tone rhythmic sequence or a continuous tone (black) cued a target tone that was either rhythmically aligned with the cue or randomly timed (red). Participants indicated by button press whether the target tone had an increasing or decreasing pitch. All combinations of (*rhythmic/random cue* x *rhythmic/random target* are shown. Solid red lines represent one presentation of the target, dashed lines show other possible timings. Note that the random target could be presented at any point during the target period.

The target was preceded by a temporal auditory cue, which could be either rhythmic or random (i.e., continuous). In the rhythmic-cue condition, we presented four identical tones at a particular presentation rate. These tones had a pitch frequency of 400 Hz, a duration of 40 ms, and a sample rate of 44100 Hz.

We used a Hanning taper to remove sharp edges. We normalized all tones, including target tones, to the same sound pressure level. In the random-cue condition, we presented the same tone continuously for a time duration that mirrored the rhythmic-cue window. Note that while ‘continuous cue’ might be the label that better reflects the nature of the cue, we have chosen ‘random’ such that we would have the same labels for the factors *cue* and *target* rhythmicity in this 2×2 design (see below).

The timing of the target presentation could similarly be either rhythmic or random. In the rhythmic-target condition, we presented the target tones either in-phase (80% of trials) or out-of-phase (20%) with respect to the preceding cue rhythm, within a window of at most four cycles, i.e., in-phase targets could occur 1, 2, 3, or 4 cycles after the cue, out-of-phase targets could occur 1.5, 2.5, or 3.5 cycles after the cue. In the random-target condition, we drew the timing of the tone from a flat probability distribution, with the full window matching that of the rhythmic-target condition. We instructed participants to respond as fast as possible via a button press.

In all three experiments, we used multiple cue frequencies, represented as their inverse or *period*, that is, the duration of one cycle. For trials with rhythmic cues, this meant manipulating the period of the tone stream; for trials with continuous cues, this meant simply manipulating the total duration of the cue. Similarly, we determined the timing of the target presentation according to that trial’s period.

Note that on rhythmic cue-rhythmic target trials, the rhythmic cue provides explicit temporal information with regard to target timing, whereas on rhythmic cue-random target trials the rhythm does not provide information beyond the length of the full window in which the target can occur. In both cases, the participant could form an (automatic) rhythmic prediction, but only in the former is it helpful for the task. On random cue-rhythmic target trials, the cue provides implicit temporal information, and cue offset can be used to predict the timing of the implicit rhythm (that is, if the participant has learned the rhythmic target-structure over the course of a block), whereas on random cue-random target trials there is no temporal information available beyond the full target window length. Whether these two conditions differ in terms of temporal predictions depends on whether the participant picks up on the implicit statistics of the task.

### Experimental protocol

Experiment I consisted of a within-subject 2×2 design with factors cue (rhythmic vs. random) and target (rhythmic vs. random), i.e., all participants performed all combinations of rhythmic/random cue/target conditions. Additionally, we used three different periods (500, 600, and 700 ms, corresponding to 2.0,

∼1.6, and ∼1.4 Hz, respectively) for the rhythmic conditions, and corresponding window lengths for the random conditions. Participants performed 12 blocks of 60 trials each, with fixed condition (i.e., rhythmic-rhythmic, rhythmic-random, random-rhythmic, random-random) and period (i.e., 500, 600, and 700 ms) per block.

Experiment II consisted of a between-subject design in which each participant performed only one of the four task conditions. We used three different periods for each participant (400, 600, and 900 ms, corresponding to 2.5, ∼1.6, and ∼1.1 Hz, respectively). Participants performed nine blocks of 60 trials each, with fixed period per block.

Experiment III consisted of only the rhythmic cue-rhythmic target condition. We used ten different periods in order to determine frequency specificity of potential temporal facilitation effects (100, 120, 150, 200, 250, 400, 600, 800, 1000, and 1250 ms, corresponding to 10, ∼8.3, ∼6.6, 5, 4, 2.5, ∼1.6, 1.25, 1, and 0.8 Hz, respectively). Participants performed ten blocks of 60 trials each, with randomized period across trials per block.

### Data analysis and statistics

We analyzed behavioral performance in terms of accuracy and reaction time (RT) and excluded participants with accuracy scores lower than 55% (see section 2.1 Participants). We included trials with rhythmic targets occurring out-of-phase (20%) when addressing whether target phase influenced performance but removed these out-of-phase trials from the data for all other analyses.

We then normalized RT per participant (for raw values, see Supplementary Figure 1) by dividing single-trial RTs by the participant’s mean RT and removed outlier trials outside the boundaries of Tukey fences (average excluded trials per participant; exp I: 36/480; exp II: 25/540; exp III: 28/480). Next, we equalized trial numbers across conditions by randomly omitting trials and excluded participants with fewer than five trials in any condition (resulting in one participant removed from experiment I).

Finally, on the remaining data, we calculated accuracy (% correct trials) per condition, then removed incorrect trials (average incorrect trials per participant; exp I: 156/480; exp II: 76/540; exp III: 73/480) and calculated mean RT per condition. The minimum, maximum, and mean number of trials that went into each of these contrasts are reported in Supplementary table 1.

For experiment I, we computed classical and Bayesian repeated measures ANOVAs to estimate differences in accuracy and RT using the factors ***cue rhythmicity*** (*rhythmic* vs. *random*), ***target rhythmicity*** (*rhythmic* vs. *random*), and ***period*** (the different cue frequencies, represented as their inverse). For experiment II we did the same but with *cue rhythmicity* and *target rhythmicity* conditions as between-subject factors, and *period* as within-subject factors. For experiment III, there was only the within-subjects factor *period*. We applied Greenhouse-Geisser correction whenever the assumption of sphericity was violated.

To estimate accuracy and RT differences in ***target phase*** (*in-phase* vs. *out-of-phase*) and whether they interacted with *cue rhythmicity*, we computed separate classical and Bayesian repeated measures ANOVAs with these two factors in experiments I and II, and a t-test to contrast in-vs out-of-phase trials in experiment III.

Across the three experiments, we modelled the hazard rate as a linear increase in accuracy, and a linear decrease in RT, across the seven possible target latencies (four in-phase latencies and 3 out-of-phase latencies). For the conditions with random target latencies, we binned the latencies into seven bins corresponding to the seven rhythmic target latencies. We pooled the data from the different conditions together and extracted, for each participant, deviations from the hazard rate predictions at the different latencies (Fiebelkorn et al., 2013; Spaak et al., 2014). We then analyzed those data in the same way as reported above.

## Results

### No benefit of rhythmicity or in-phase target presentation

First, we investigated whether there was any benefit of rhythmic cues and/or targets (vs. random ones) on behavioral performance (RT and accuracy). We manipulated the rhythmicity of the cues and targets in experiments I (within-subjects) & II (between-subjects), and found the same pattern of results in both experiments (Figure 2).

**Fig. 2:**
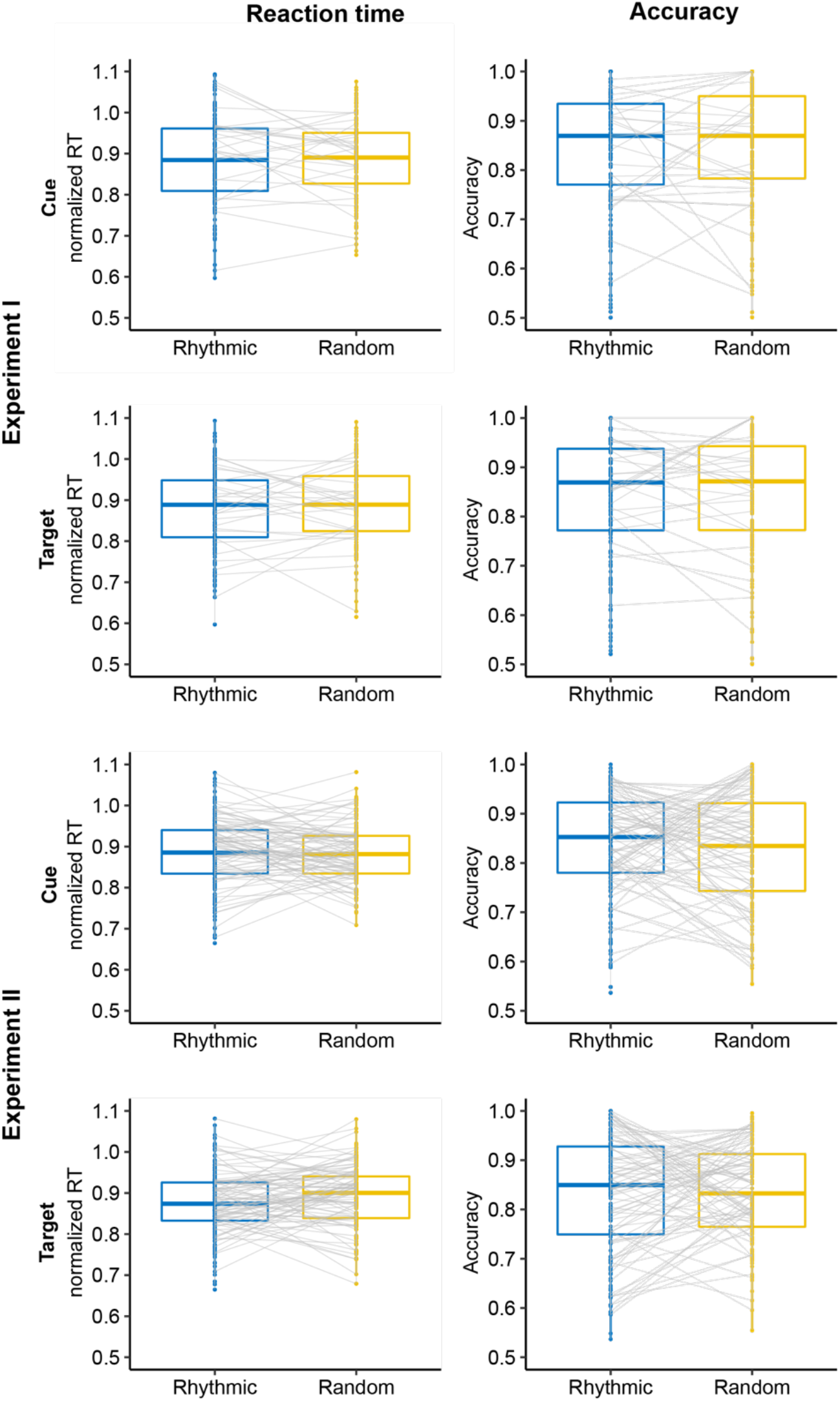
Effect of cue and target rhythmicity. In both experiment I (within-subjects; top half) and experiment II (between-subjects; bottom half), participants were neither faster (left) nor more accurate (right) in responding to the target when the cue was rhythmic (vs. random) or when the target was rhythmic (vs. random).

**Fig. 3:**
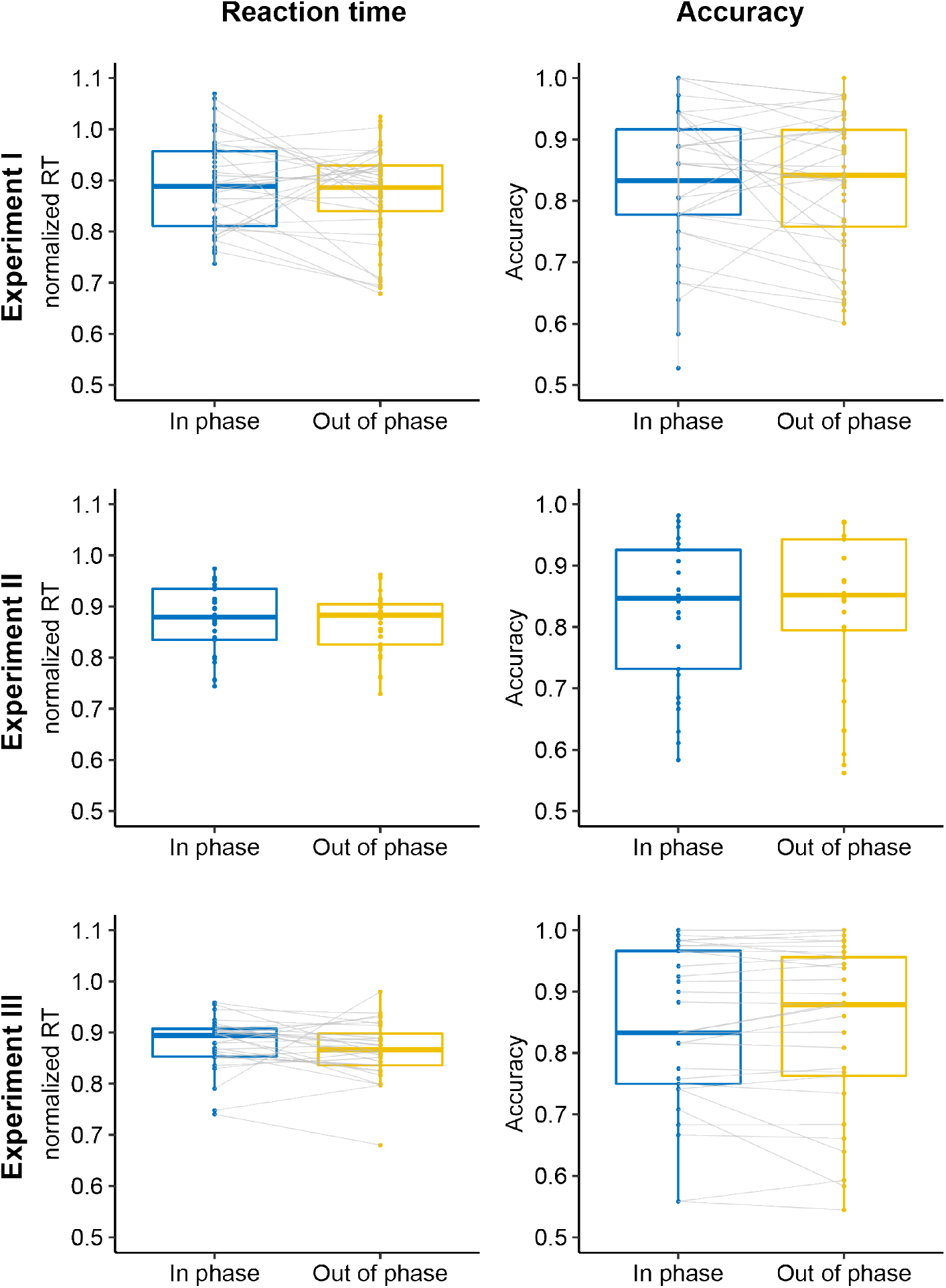
Effect of phase. Across all three experiments (top, middle, and bottom panels), participants were neither faster (left) nor more accurate (right) in responding to a target occurring in-phase (vs. out-of-phase) with the cue stream.

In experiment I, whether the cue was rhythmic or random had no effect on RT (*F(1,29)* = 0.86, *p* = .362, *BF*_10_ = 0.09) nor on accuracy (*F(1,29)* = 2.57, *p* = .12, *BF*_10_ = 0.172), and whether the target was rhythmic or random also had no effect on RT nor on accuracy (RT: *F(1,29)* = 0.22, *p* = .64, *BF*_*10*_ = 0.054; accuracy: *F(1,29)* = 0.71, *p* = .4, *BF*_*10*_ = 0.058). Similarly, in experiment II, the cue rhythmicity had no effect on RT (*F(1,110)* = 8.1e-5, *p* =.92, *BF*_10_ = 0.58) nor on accuracy (*F(1,110)* = 0.36, *p* = .55, *BF*_10_ = 0.22), and target rhythmicity also had no effect on RT (*F(1,110)* =2.33, *p* = .13, *BF*_*10*_ = 0.38) nor on accuracy (F*(1,113)* = 0.15, *p* = .69, *BF*_*10*_ = 0.19).

Moreover, contrary to our expectations, none of the interactions showed a significant effect. This included the interaction of interest in the context of entrainment: i.e., between rhythmicity of cue and target in experiment I (RT: *F(1,29)* = 0.19, *p* = .66, *BF*_*10*_ = 0.01; accuracy: *F(1,29)* = 0.04, *p* = .83, *BF*_*10*_ = 0.01) and experiment II (RT: F(1,110) = 3.12, p = .08, B_10_ = 0.54; accuracy: F(1,110) = 0.12, p = .72, B_10_ = 0.12). In other words, people were not better at discriminating a target tone occurring at a predictable time point after a rhythmic cue, compared to when the target tone occurred at a random time point after a random cue.

It could be argued that potential rhythmicity effects might only be observed at early target latencies, for example at the first possible target position, as rhythmicity effects might fade or vanish at later positions. To account for this, we repeated the above analyses only including targets occurring at the first post-cue position, and still found no effect of cue rhythmicity (RT: F(1,27) = 2.22, p = .147, BF_10_ = 0.31; accuracy: F(1,27) = 0.004, p = .948, B_10_ = 0.16) nor target rhythmicity (RT: F(1,27) = 0.905, p = .35, BF_10_ = 0.23; accuracy: F(1,27) = 1.94, p = .17, B_10_ =0.35) in experiment I nor in experiment II (cue RT: F(1,110) = 2.18, p = .14, B_10_ =0.37; target RT: F(1,110) = 2.54, p = .113, B_10_ =0.442; cue accuracy: F(1,110) = 0.004, p = .948, B_10_ = 0.138; target accuracy: F(1,110) = 0.01, p =.92, B_10_ = 0.139). Note that there was a cue x target rhythmicity interaction for accuracy in experiment I (F(1,27) = 5.3, p = .029, B_10_ = 0.17), but none of the post-hoc contrasts were significant.

Next, we asked whether targets occurring in-phase with a rhythmic cue were better discriminated compared to targets occurring out-of-phase, and found no evidence for such an effect on RT (experiment I: F(1,28) = 0.7, p = .41, BF_10_ = 0.85; experiment II F(1,57) = 2, p = .16, BF_10_ = 0.34; experiment III: t(28) = 1.24, p = .22; BF_10_ = 0.39) nor accuracy (experiment I: F(1,28) = 0.11, p = .74, BF_10_ = 1.52; experiment II F(1,57) = 0.03, p = .85, BF_10_ = 0.14; experiment III: t(28) = 0.35, p = .72 BF_10_ = 0.2). This did not depend on whether the cue was rhythmic or random i.e., no significant cue-by-phase interaction for RT (experiment I: F(1,28) = 0.06, p = .8, BF_10_ = 0.07; experiment II: F(1,57) = 0.23, p = .63, BF_10_ = 0.13) nor accuracy (experiment I: F(1,28) = 1.82, p = .19, BF_10_ = 0.17; experiment II: F(1,57) = 0.12, p = .72, BF_10_ = 0.064) Finally, it could be argued that phase effects are only observable for particular, individually preferred frequencies, and that including multiple rhythms dilutes this effect at the group level. To account for this concern, we repeated this analysis for experiment III, only including individually preferred rhythms (defined as the frequency with the highest accuracy) and still found no effect of in-vs. out-of-phase target presentation (*t*(28) = −0.06, *p* = .954, BF_10_ = 0.21).

### Better performance for later-occurring targets

We then asked whether the duration of the time window between cue offset and target onset (the cue-target delay) had an influence on behavioral performance. The reasoning is that as time after cue offset increases, so does the probability of the target occurring. We would expect this hazard rate effect to lead to better performance on later occurring targets.

To disentangle this non-rhythmic temporal expectation effect from any possible rhythmic entrainment effects, we modelled the hazard rate for each participant and extracted deviations from the predicted outcomes. We modelled accuracy at each target position as linearly increasing from the first to the last position, and reaction time as linearly decreasing. On average, the accuracy trend had a positive slope (experiment I = .014; experiment II = .012; experiment III = .007) and the RT trend had a negative slope (experiment I = −0.017; experiment II = −0.014; experiment III = −0.008). These slopes were significantly different from zero based on one-sampled t-tests (all p < .001, results in table 1), suggesting that participants indeed performed better for later-occurring targets (Figure 4).

**Table 1:** Hazard rate effects. Results from the one-sample t-tests (against zero) of the estimated slopes of accuracy and RT trends in all experiments. Generally, accuracy trend had a positive slope and the RT trend had a negative slope.

| Experiment | Measure | t-value | df | p-value |
| --- | --- | --- | --- | --- |
| Experiment I | Accuracy | 6.05 | 31 | < .0001 |
|  | RT | -3.37 | 31 | .002 |
| Experiment II | Accuracy | 9.34 | 118 | < .0001 |
|  | RT | -7.79 | 118 | < .0001 |
| Experiment III | Accuracy | 4.00 | 29 | < .0001 |
|  | RT | -3.95 | 29 | < .0001 |

**Fig. 4:**
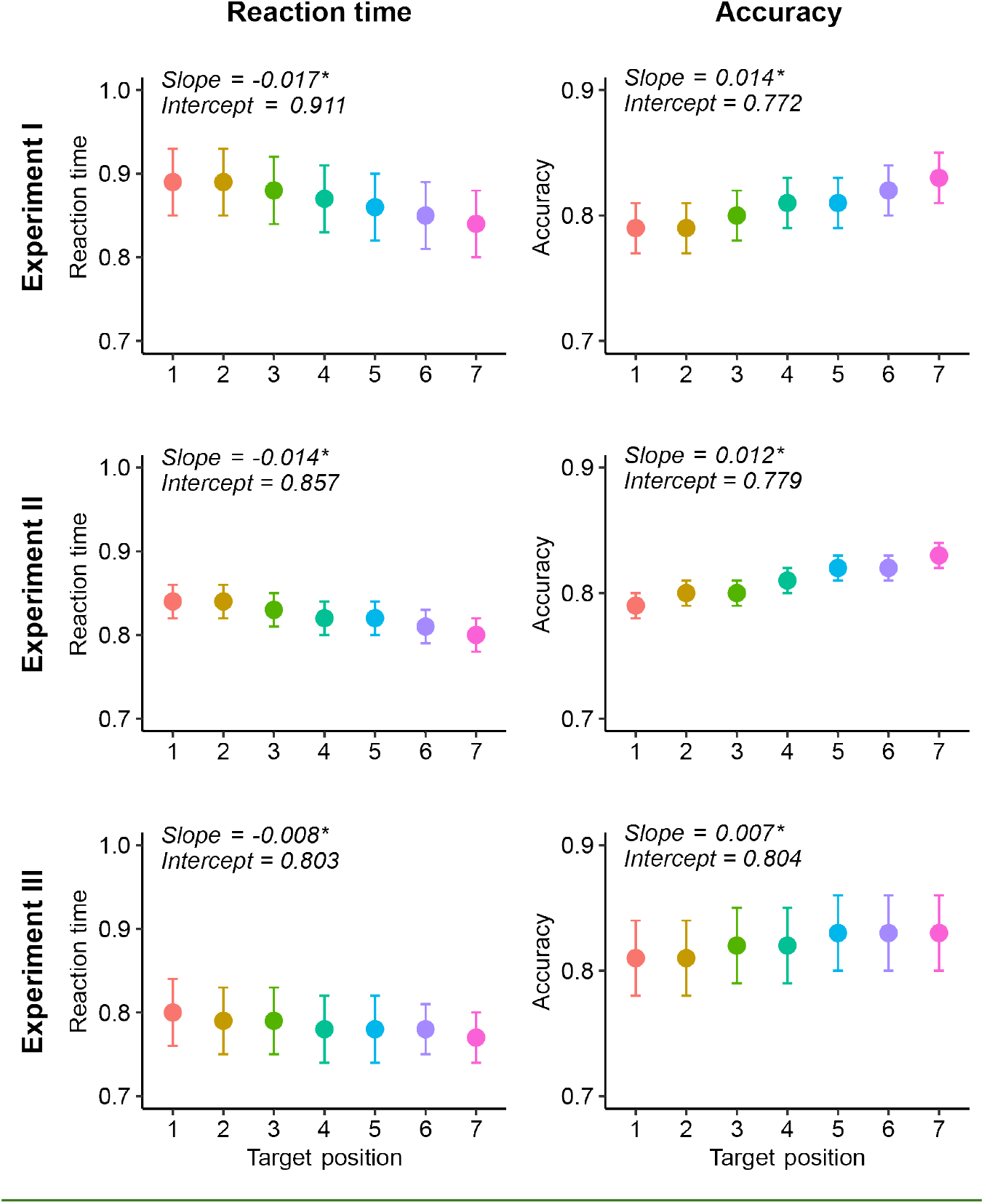
Hazard rate effect. In all three experiments, we modelled the hazard rate effect across the seven possible target onsets. Left panels: participants responded faster, the later the target onset was. Right panels: participants responded more accurately, the later the target onset. Asterisk indicates slope is significantly higher or lower than zero.

We then repeated the analyses reported in the previous section, but with the detrended values extracted after modelling the hazard rate. These results largely conformed with the ones reported above. That is, after accounting for the hazard rate, cue and target rhythmicity and whether the target occurred in or out of phase still had no influence on performance (Supplementary Tables 2-7). In sum, this analysis suggests that despite non-rhythmic temporal expectation effects (described by hazard rate) being present in our data, rhythmic entrainment effects were not.

### Faster responses following faster rhythmic cues

Our last question whether some cueing rhythms were more behaviorally beneficial than others (Figure 5). Across all three experiments, we observed a remarkably robust speeding up of RT with faster cues (exp I: *F(2,58)* = 15.01, *p* < .0001, *BF*_*10*_ > 100; exp II: *F(2,220)* = 80.42, *p* < .001, *BF*_*10*_ > 100; exp III: *F(9,243)* = 34.87, *p* < .001, *BF*_*10*_ > 100). This effect was most evident in experiment III, which included ten different periods rather than three, and where the cues were exclusively rhythmic and varied trial-wise (in frequency) rather than block-wise. Accuracy also increased with faster cues in experiment II (*F(2,220)* = 6.6, *p* = .002, *BF*_*10*_ = 4.46), but not in experiments I and III (exp I: *F(2,58)* = .1, *p* = .9, *BF*_*10*_ = .014; exp III: F*(9,243)* = .93, *p* = .49, *BF*_*10*_ = .018), suggesting the RT effect is not necessarily reflective of a speed-accuracy trade-off.

**Fig. 5:**
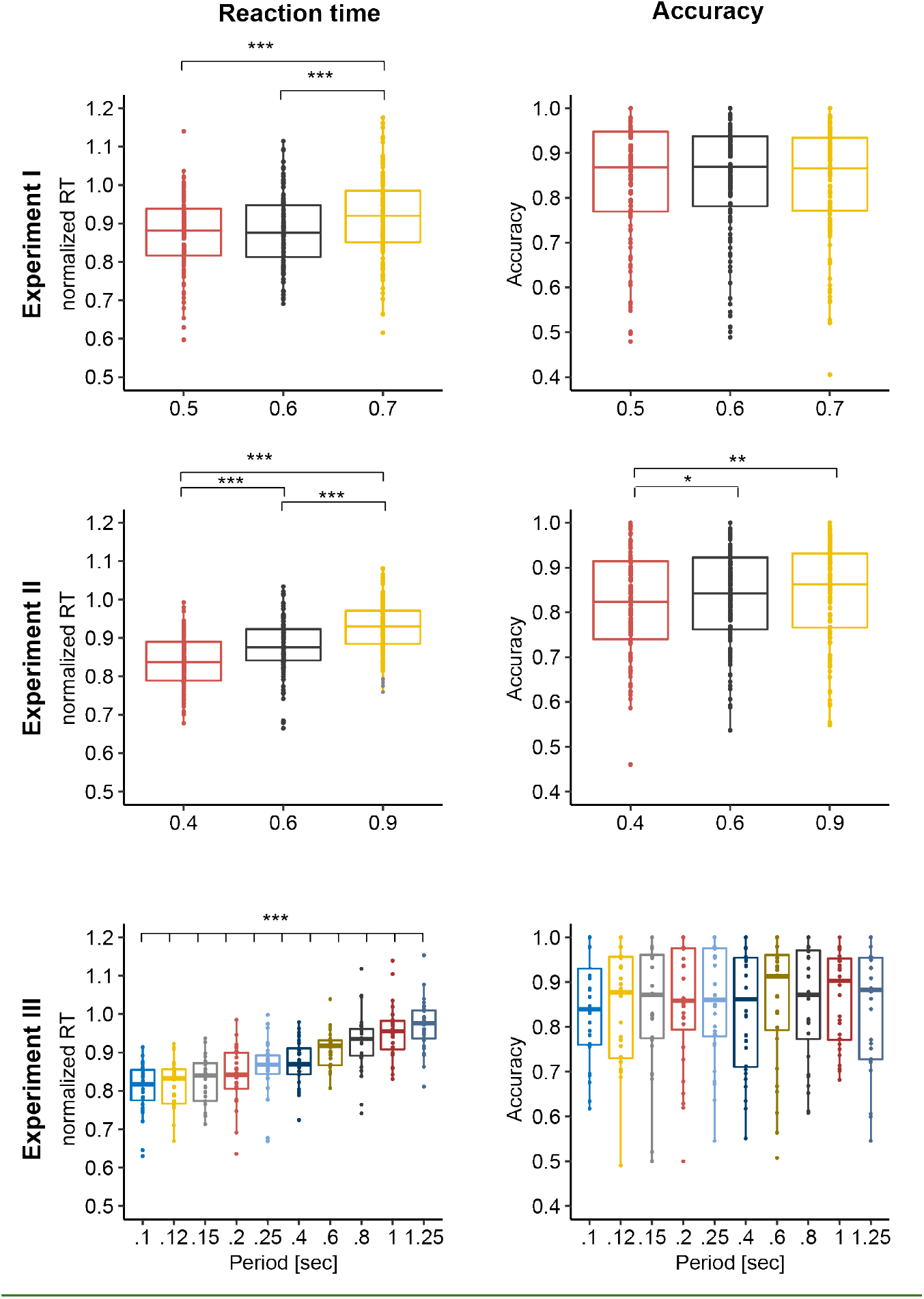
Effect of period. Left panels: In all three experiments, participants were faster to respond on trials with faster cueing frequencies (shorter *period*). Right panels: In experiment II (middle) but not experiments I and III (top and bottom), participants were more accurate in responding on trials with slower cueing frequencies (longer *period*). *p<0.05; **p<.01; ***p<.001.

In experiments I and II, where cues and targets could be rhythmic or random, we then asked whether the rhythmicity of cues and targets interacted with the RT effect of period reported here (Figure 6). In experiment I, the effect of faster cues on RT depended neither on the rhythmicity of the cue (*F(2,58)* =

**Fig. 6:**
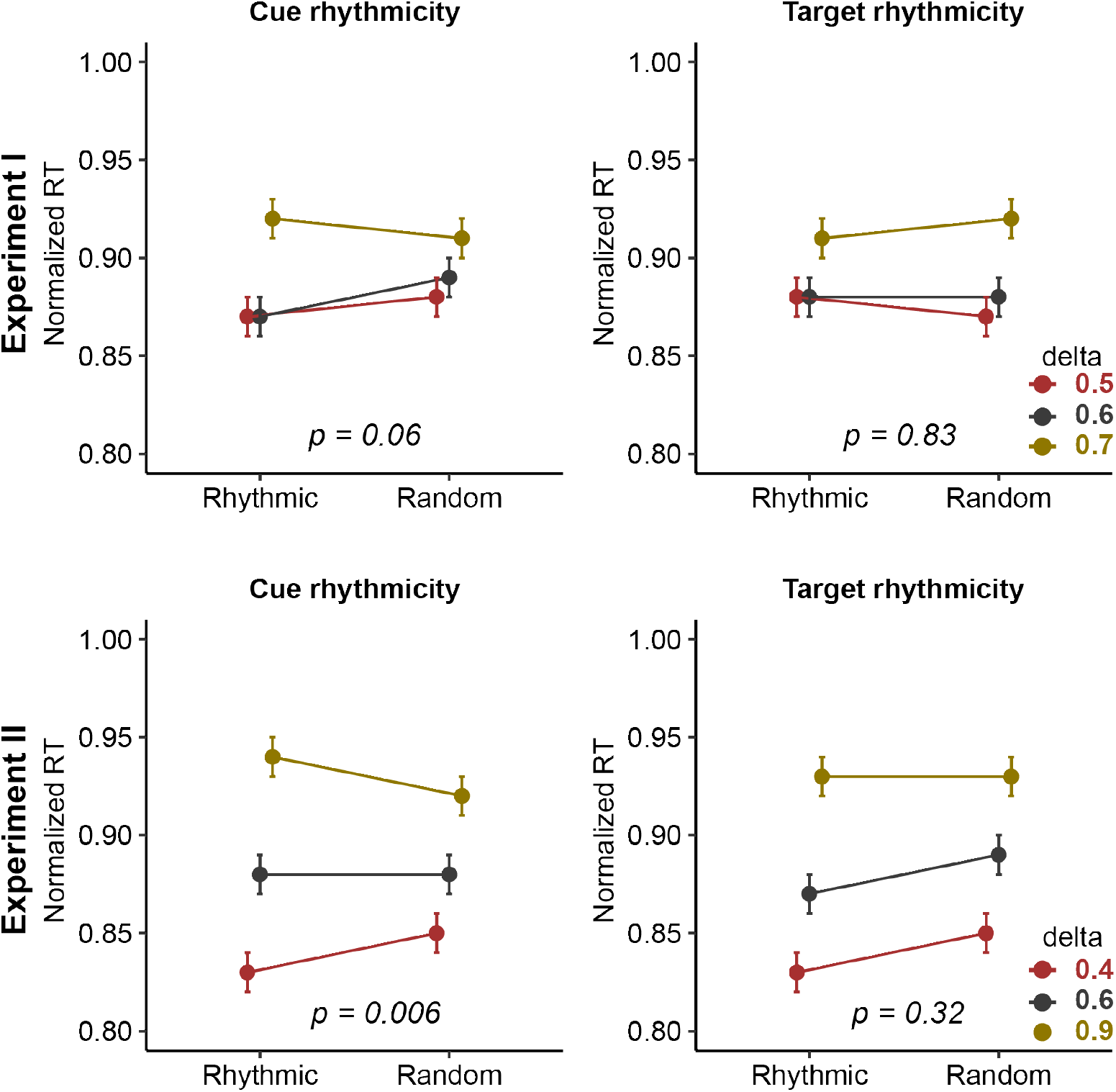
Interaction between period and cue/target properties. Left panels: In experiment I and II the speeding up of responses with faster cueing frequencies (shorter *period*) was more prominent when the cue was rhythmic (compared to random). Middle panels: In both experiments there was no interaction between the cueing frequencies and the target rhythmicity. Left panels: In both experiments there was no interaction between the cueing frequencies and the target timing. Error bars represent standard error of mean.

3.05, *p* = .06, *BF*_*10*_ = 0.1), nor that of the target (*F(2,58)*= 0.17, *p* = .83, *BF*_*10*_ = 0.01). In experiment II, the RT effect of period did not depend on target rhythmicity (*F(2,220)*= 1.014, *p* = .32, *BF*_*10*_ = .54), but it did depend on cue rhythmicity (*F(2,220)* = 5.16, *p* = .006, *BF*_*10*_ = 2.35), such that rhythmic cues led to more RT speed-up. We could not test this in experiment III because we only used rhythmic cues. Finally, the RT effect of period held robustly after accounting for the the hazard rate effect (Supplementary tables 2-4). Overall, the speeding up of RT following faster cues seems to benefit weakly from rhythmic cueing.

## Discussion

In a series of three experiments, we found no behavioral benefit of rhythmic cueing, compared to random cueing (i.e., a non-rhythmic continuous tone), neither on a within-subjects level (experiment I) nor on a between-subjects level (experiment II). We also found no behavioral advantage for targets appearing at a rhythmically consistent timing, compared to those appearing at a random timing. In addition, we found no behavioral benefit for targets occurring in-phase with rhythmic cues, compared to those occurring out-of-phase. However, we found that shortening the duration of the cue — that is, speeding it up — consistently resulted in faster reaction times.

The idea of neural entrainment as a mechanism to facilitate sensory processing rests on the assumption that neural oscillations reflect rhythmic phases of high and low neural excitability that coincide with phases of good and bad perceptual performance, respectively. This is supported by evidence that such phase effects occur spontaneously, i.e., without exposure to an external rhythm (e.g., Busch et al., 2009; Mathewson et al., 2009). Within the entrainment framework, it is then thought that these internal phases can be adjusted to external rhythms, potentially providing a mechanism for temporal prediction. The influence of external rhythms on perception and subsequent behavior has been tentatively shown (e.g. Jones et al., 2002, 2006), but these results are now being scrutinized by the field, for example in the current special issue (also see Haegens & Golumbic, 2018 for review). From an electrophysiological point of view, it has been difficult to show that neural oscillatory phase entrains to external rhythms (e.g. Wilsch et al., 2020), as an observed “entrained” brain rhythm is difficult to disentangle from a series of evoked responses, a series of top-down predictions, or simple resonance (Haegens, 2020; Helfrich et al., 2019; Obleser & Kayser, 2019). Since we did not collect electrophysiological data in our studies, we will restrict our discussion to the behavioral aspect of entrainment.

If the assumptions of entrainment are met, one would expect the entrained neural oscillations (and hence the concomitant behavioral benefit) to persist after the external rhythm stops (Lakatos et al., 2008). That rhythmicity in input streams offers perceptual and behavioral advantages has been shown repeatedly (Henry & Obleser, 2012; Jones et al., 2002, 2006; Rohenkohl & Nobre, 2011; Rohenkohl et al., 2011), however, most of these studies report these advantages when targets occur *within* rhythmic streams, with very few reporting advantages persisting after the stream stops. To the best of our knowledge, the few studies that have reported a persistent advantage have relied on relatively low numbers of participants (e.g. Farahbod et al., 2020; Hickok et al., 2015; five participants each; Mathewson et al., 2010, 2012; 13-16 participants each) and did not explicitly test for temporal predictions (i.e., rhythmic cues were uninformative). In our current study we tested whether rhythmicity in an auditory cue stream influences the discrimination of a target occurring after offset of the stream and found no such evidence.

There are several possible explanations for these discrepancies: first, the nature of the task (detection versus discrimination) could play a role in limiting the behavioral facilitation of neural entrainment (Bauer et al., 2015). Rhythmic facilitation has been observed in demanding detection tasks, for example where near-threshold targets are embedded in noise (e.g. Ten Oever et al., 2017), and arguably more precise temporal predictions are needed. Though our stimuli were short-lived and the discrimination task fairly demanding, it is possible that rhythmic facilitation effects are only (or mostly) relevant in paradigms where time is a more critical factor, such as in speeded tasks, time estimation, and near-threshold detection. However, if that is the case, it would argue against entrainment as an automatic, bottom-up effect (Haegens & Golumbic, 2018). It is also possible that rhythmicity only impacts early-occurring targets as the effect fades over time, but we found no evidence for this in our data. Furthermore, it has been suggested that temporal expectation effects mostly boost other forms of attention and prediction (Rohenkohl et al., 2014; Morillon et al., 2016), i.e., perhaps by themselves the effects are too weak to detect in most scenarios.

Other possibilities are that four cue tones were not enough to build temporal expectations (though see Breska & Deouell, 2014), or that the variability in the rhythms used made it harder to build expectations and predict cue onset (though note we used a blocked design in experiments I and II). The predictability of rhythmic cues was previously shown to correlate with the degree of phase alignment of brain oscillations (Stefanics et al., 2010; but see Breska & Deouell, 2017), something we cannot assess here as we did not collect electrophysiological data. Furthermore, one could posit that if participants are exposed to both rhythmic and random cues in a single experiment, a less cognitively-demanding strategy is to entirely ignore the cues (both rhythmic and random) as they provide no perceived behavioral benefit. However, our experiment III was designed with cue rhythmicity as a between-subject factor to avoid such a carry-over effect, and nevertheless we found no effect of cue rhythmicity.

Further, there might be interindividual variability in preferred frequency (and phase) on which such behavioral benefits depend (Zoefel et al., 2018), so the use of one frequency (and phase) for all participants might not lead to an observable effect at the group level. In all our experiments we used multiple frequencies (three in experiment I-II and ten in experiment III) and still did not find an impact of cue rhythmicity for any of the frequencies, not even when taking into account interindividual variability in preferred frequencies. Another possible source of interindividual variability is that different people might have different sensitivity to entrainment (Assaneo et al., 2019). If this were the case in our experiments, we would expect the individual-level effects to be bi-modally distributed, but the distributions of effects do not suggest this (Supplementary Figures 2-4). Finally, musical ability and training have been suggested as factors contributing to interindividual differences in entrainment, with musicians being better entrainers (Doelling & Poeppel, 2015). However, if true, would render entrainment less likely as a candidate mechanism for general temporal prediction.

Another question we aimed to address was whether different cueing frequencies had different effects on behavior either due to the mechanistic roles ascribed to these frequencies: ramping up activity at slower frequencies (delta to theta) could lead to behavioral facilitation, as these low-frequency rhythms are thought to play a role in sensory sampling (Fiebelkorn & Kastner, 2019; VanRullen, 2016), while a similar increase in higher frequencies (especially alpha) could lead to a behavioral cost, as these oscillations are thought to play a role in functional inhibition (Klimesch et al., 2007). We found no evidence for certain frequencies inducing differential behavioral effects. Instead, we found that faster cues led to faster responses, a robust effect observed in all three experiments but most strikingly in experiment III, which was designed to address this question on a trial-by-trial level.

In paradigms with varying cue-target delays, RTs are usually faster and accuracy scores higher on trials with long delays, as uncertainty of target timing decreases the later the target occurs (hazard rate effect; Näätänen, 1971). This hazard rate effect was present in our data but did not confound any of our results. In fact, all our results held even after accounting for the hazard rate. In our experiment, faster cues were followed by earlier-occurring targets on average, so based on the hazard rate prediction, one would expect responses to faster cues to be slower, but we found them to be faster. Taken together, these findings suggest that hazard rate and the observed RT speed-up with faster cues were dissociated in our data.

Our results can tentatively be explained with the notion of covert active sensing: i.e., the motor system actively coordinates the sensory system to adjust to the current environment (Schroeder et al., 2010). As a result, a faster external rhythm might increase the communication rate between the sensory and motor cortices. A faster communication rate in turn gives a faster response on average, since input can be sampled earlier in time. This interpretation is particularly supported by experiment III, where different frequencies were randomized across trials (i.e., not blocked as in the first two experiments), suggesting this is a rapidly adaptive mechanism suitable for real-life situations with varying temporal (ir-)regularities. Future work should further address this potential mechanism on the neural level, and, more generally, the role of the motor system in (rhythmic) temporal prediction (Balasubramaniam et al., 2021; Cannon & Patel, 2020).

Finally, we found that faster responses followed faster cues, particularly when the cues were rhythmic. The RT speed-up depended on cue rhythmicity in experiment II and showed a similar trend in experiment I (note that we used smaller differences between periods in experiment I). We do not offer conclusive evidence as to whether this RT effect is exclusive to rhythmic contexts, but rhythmicity has been shown to improve response readiness (Morillon et al., 2016). Future work should address this question by manipulating cue rhythmicity on a trial-by-trial basis, in addition to independently varying the speed and duration of the cue. That is, in our current design, length of cue and target window scaled with the frequency of the cued rhythm (since we used a fixed number of cycles). Separately manipulating these factors would allow disentangling effects driven by the cue rhythm per se, versus effects driven by the task rhythm. This would provide more insight into whether this speed-up is specific to micro (i.e., within trial) rhythmic contexts or reflects a more general adaptation to faster macro (i.e., across trials) rhythms.

## Acknowledgements

We would like to thank Aarti Ramchandran and Camille Gret for assistance with data collection, and Sanne ten Oever for thoughtful comments on an earlier version of the manuscript. This work was supported by NWO grants Veni 451-14-027 and 016.Vidi.185.137 to SH.

## Data Accessibility

The data that support the findings of this study and the code that was used to analyze it will be publicly available after publication, on the OSF page https://osf.io/spt24/.

## Author contributions

HE, SH, ER, & AW conceived and designed the study. IB-M, JE, WML, & DO collected the data. HE, WML, DO, & ER analyzed the data. HE, SH, WML, DO, & ER wrote the manuscript.

